# An updated global COI barcode reference data set for Fall Armyworm (*Spodoptera frugiperda*) and first record of this species in Bhutan

**DOI:** 10.1101/2020.05.17.100883

**Authors:** Kiran Mahat, Andrew Mitchell, Tshelthrim Zangpo

## Abstract

We report the first detection of Fall Armyworm (FAW), *Spodoptera frugiperda* (Smith, 1797), in Bhutan. FAW feeds on more than 300 plant species and is a serious pest of many. It has been spreading through Africa since 2016 and Asia since 2018. In Bhutan, this species was first detected in maize fields in the western part of the country in September 2019 and subsequently found infesting maize crop in southern parts of the country in December 2019 and April 2020. Using morphological and molecular techniques the presence of the first invading populations of *S. frugiperda* in Bhutan is confirmed through this study. We present an updated reference DNA barcode data set for FAW comprising 374 sequences, which can be used to reliably identify this serious pest species, and discuss some of the reasons why such compiled reference data sets are necessary, despite the publicly availability of the underlying data. We also report on a second armyworm species, the Northern Armyworm, *Mythimna separata* (Walker, 1865), in rice, maize and other crops in eighteen districts of Bhutan.

## Introduction

The Fall Armyworm (FAW), *Spodoptera frugiperda* (Smith, 1797), is an invasive pest species native to the Americas that has been spreading rapidly through Africa since 2016, in Asia since 2018, and in 2020 arrived in Australia via Papua New Guinea (Goergen et al., 2016; Kalleshwaraswamy et al., 2018; CABI, 2020). FAW feeds on more than 300 species including grains (rice, maize, sorghum), other grasses, many non-grain crops (e.g., soybean, peanut, potato, sweet potato, spinach, tomato, etc.) and can cause devastating losses (Day et al., 2017; Montezano et al., 2018). Like many pests, it is the larval stage that causes the damage, yet larvae are difficult to identify. Although larval identification resources have been developed for this species (Gilligan and Passoa, 2014), DNA-based methods provide an independent and universal identification method and are applicable to all life stages.

DNA barcoding is an increasingly useful tool for identifying arthropod plant pests (Ashfaq and Hebert, 2016) and moths of quarantine concern in particular (e.g., Mitchell and Gopurenko, 2016; Lee et al., 2019). Due to the threat that pest insect incursions pose to agricultural crops, there is a need for comprehensive and well-curated databases of DNA barcode sequences to identify intercepted specimens. Unfortunately, the indiscriminate use of the BOLD and GenBank databases is ill-advised, since both databases contain examples of misidentified sequences, which may lead to confusion in assigning a species name to an unknown sample (Lee et al., 2019). The BOLD platform at least offers the advantage that it is a curated database, meaning that such errors can be corrected by subsequent studies. Such errors were encountered during the course of routine diagnostics work aiming to identify armyworms and are corrected in this study. We report on the first detection of Fall Armyworm, *Spodoptera frugiperda* (Smith, 1797), in Bhutan, and on infestations of Northern Armyworm, *Mythimna separata* (Walker, 1865).

## Material and Methods

### Sample collection

The FAW samples were collected in September 2019 from a maize field in Guma, while the Northern Armyworm samples were collected in May 2013 from Shengana (Table 1). Both sites are in Punakha Province in western Bhutan.

**Table 1.** Sample collection and DNA sequence accessions.

| Sample ID | Species | Collectors | Collection Date | Country | Locality | GenBank Accession |
| --- | --- | --- | --- | --- | --- | --- |
| am14016 | *Spodoptera frugiperda* | Kiran Mahat, Tshelthrim Zangpo | 17.09.2019 | Bhutan | Guma | MT324067 |
| am14017 | *Spodoptera frugiperda* | Kiran Mahat, Tshelthrim Zangpo | 17.09.2019 | Bhutan | Guma | MT324066 |
| am14018 | *Spodoptera frugiperda* | Kiran Mahat, Tshelthrim Zangpo | 17.09.2019 | Bhutan | Guma | MT324065 |
| am11901 | *Mythimna separata* | Kiran Mahat | 10.05.2013 | Bhutan | Shengana | MT324064 |
| am11902 | *Mythimna separata* | Kiran Mahat | 10.05.2013 | Bhutan | Shengana | MT324063 |

### Sample identification

Specimens were identified by dissecting male genitalia and comparing them with the literature (Pogue, 2002, for *Spodoptera frugiperda*; Franclemont, 1951, for *Mythimna separata*).

### DNA barcoding

A single leg was removed from adult moths for DNA extraction. Forceps were wiped with lab tissue, dipped in 80% ethanol and flame-sterilized between samples. DNA was extracted using a Bioline Isolate II Genomic DNA Isolation Kit (Bioline, Eveleigh, Australia) following the manufacturers’ instructions, except that the final elution volume was adjusted to 70 µL. PCR amplification used the protocols and primers described in Mitchell (2015), with a MyTaq HS Red Mix PCR kit (Bioline, Eveleigh, Australia). Each 20 μL reaction contained 5 μL of MilliQ water, 10μL of MyTaq HS Red Mix, 0.4μL of 50mM MgCl_2_, 0.8μL each of forward and reverse primers at 5 μM, and 3μL of DNA template. PCR product purification and bidirectional Sanger sequencing was performed by Macrogen Inc. (Seoul, South Korea). Chromatograms were edited and consensus sequences generated using Geneious 10.2.6 (Kearse et al., 2012).

DNA sequences were submitted to the BOLD and GenBank databases. Specimen collection data and sequence accession numbers are provided in Table 1.

The five sequences we derived in this study were searched against both the BOLD (All Barcode Records) and GenBank databases on 17 October 2019 to obtain confirmation of their identity. The *Spodoptera frugiperda* sequences were subjected to further analyses because 1) the searches gave inconclusive results, and 2) we hoped to ascertain the geographic origin of the population recently detected in Bhutan.

We downloaded COI sequences for *Spodoptera frugiperda* from both BOLD and GenBank on 17 October 2019. The sequences were renamed using FaBox 1.5 (Villesen, 2007) so that the name started with the GenBank accession number (or the prefix “BOLD-” for sequences not in GenBank), followed by the sample’s country of origin and host plant, where this information was provided, or could be ascertained from the published paper associated with the sequences. In some instances, the country of origin or host plant was implied in the paper but not overtly stated, and in such cases the information is provided in brackets in the sequence title. Duplicate sequences (those on both GenBank and BOLD) were deleted, and remaining sequences were aligned in Geneious using the MUSCLE option. The 60 sequences KT809235–KT809294 all contain a 100 nt stretch of Ns, starting at position 595 of the alignment. It is clear from the difficulty aligning these sequences that the authors sequenced two non-overlapping fragments of COI and joined them with an arbitrary number of Ns. For all 60 sequences we deleted data from the start of these Ns to the end of the sequence, leaving 556 nt of aligned sequence data. The resulting alignment was cropped to a length of 658 bp, the standard barcode region for insects. The 16 sequences that were less than 500 bp in length were then deleted.

FastTree 2.1.5 (Price *et al.*, 2010) was used to generate approximately maximum-likelihood trees. A first analysis was used to identify publicly available sequences which had likely been misidentified to species. Such sequences were then searched against the BOLD database to identify them, and if the conclusion was that they had been misidentified, they were excluded from the final data set. A number of sequences were identified as forming very long terminal branches in the initial tree. These sequences contained a series of substitutions in either the first 20 bases or the last 20 bases of the sequence, but showed rare or few substitutions in the rest of the sequence. In all cases these sequences were originally published in GenBank and were not associated with raw data (trace files) on BOLD, which would have allowed us to check the accuracy of the base calling. These end substitutions were therefore judged to be likely the result of poor quality data at the ends of sequences, and the mismatching terminal bases were trimmed from the alignments. Eight sequences were trimmed in this manner, and the details are provided in Table 2.

**Table 2.** End trimming of GenBank sequences.

| GenBank Accession | BOLD ProcessID | Traces on BOLD? | No. of 5'-end nucleotides trimmed | No. of 3'-end nucleotides trimmed |
| --- | --- | --- | --- | --- |
| MK295625 | GBMNA44462-19 | No | 8 |  |
| MK327538 | GBMNA44682-19 | No | 2 |  |
| MK908223 | GBMNA47805-19 | No | 2 |  |
| MK041922 | GBMNA41994-19 | No | 2 |  |
| GU094756 | PHMO358-03 | No | 15 |  |
| KT809235* | GBGL25125-19 | No |  | 16 |
| KT809240* | GBGL25130-19 | No |  | 14 |
| MH899610 | GBGL25203-19 | No |  | 12 |

\*The 60 sequences KT809235–KT809294 were all trimmed from the start of a 100 nt stretch of Ns, starting at position 595 of the alignment.

## Results

For *Spodoptera frugiperda*, three individuals were found to be misidentified (Table 3) and a fourth individual had a frameshift mutation (one base pair deletion); all four sequences were excluded from the final data set.

**Table 3.** Sequences excluded from reference data set.

| BOLD ProcessID | GenBank accession | Correct species name | BOLD species name | GenBank species name | Reason for exclusion |
| --- | --- | --- | --- | --- | --- |
| SWKFO001-19 | n/a | *Spodoptera evanida* | *Spodoptera frugiperda* | n/a | Incorrect identification. |
| MILEP712-11 | n/a | *Spodoptera evanida* | *Spodoptera frugiperda* | n/a | Incorrect identification. |
| LEMMZ129-10 | JF854742 | *Spodoptera eridania* | *Spodoptera eridania* | *Spodoptera frugiperda* | Incorrect identification. |
| GBGL25082-19 | MH794141 | *Spodoptera frugiperda* | *Spodoptera frugiperda* | *Spodoptera frugiperda* | Has 1 nt deletion at pos. 556, 10 unique base substitutions in middle, and no trace files. |
| GBMIN83714-17 | KM362176 | *Spodoptera frugiperda* | *Spodoptera frugiperda* | *Spodoptera frugiperda* | This is the original sequence that GenBank reference sequence NC_027836 (BOLD Process ID: GBMTG5891-16) is derived from, therefore this sequence is duplicated in BOLD. |

Combined with the end-trimming of eight sequences, this eliminated a number of apparent errors in sequences on BOLD and GenBank. A further 16 sequences were excluded because they were less than 500 bp in length, leaving a final COI barcode dataset of 374 sequences. This COI dataset can serve as a reference dataset for Fall Armyworm identification and is provided in FastA format on the Mendeley Data archive.

The approximate maximum likelihood tree produced by Fasttree is provided as supplementary Figure S1. In summary, the tree was deeply divided into two major clusters, corresponding to the “corn” (maize) and “rice” strains of the species, with minimum and maximum Kimura 2-parameter distances between the two clusters of 2.1–2.9%.

The “corn strain” is shown on pages 1-2 of Fig. S1 and contains 109 sequences. Sequence names comprise GenBank Accession number (or BOLD Process ID for the four BOLD sequences not on GenBank), country of collection, host plant, and “strain” which is either “Rice” or “Corn”.

All but two of the 88 host plant records for this strain are maize. The other two records are sugarcane in India, GenBank accession MK295625, for a larva in Maharashtra (Chormule et al., 2019) and MK908223 from Andhra Pradesh (Bhavani et al., 2019). Of the 109 sequences, 83 are from the Americas (Canada, USA, Puerto Rico, Mexico, Honduras, Peru, Brazil), 20 are from Africa (São Tomé and Príncipe, Ghana, Uganda, Kenya) and six are from Asia (India, China).

The “rice strain” consists of 265 sequences, from seven countries in the Americas, seven African countries and six Asian countries, and the samples were collected from four host plant species, including rice. The sequences nearest the base of the “rice strain” group include 11 samples (6 divergent haplotypes) from rice in the USA, plus samples from maize in the USA, Brazil, China and India. However, 79% of the rice strain sequences (210 sequences) are identical. This single common haplotype contains the three sequences from Bhutan specimens plus samples from the following countries (number of samples in brackets): Canada (4), USA (15), Costa Rica (17), Dominican Republic (1), Suriname (1), Brazil (10), Nigeria (4), Ghana (7), Uganda (1), Kenya (74), South Africa (12), India (31), Myanmar (1), China (25) and Vietnam (4). Host plants are recorded for 141 of these 210 samples, and all but five of them are maize; the other host plants are *Capsicum* (one sample from Dominican Republic), rice (two samples from India and one from USA) and sorghum (one sample from India).

For *M. separata*, species identity was established by dissection of male genitalia (Fig. 1), which bear the long spine on the inner margin of the cucullus, characteristic of this species (Franclemont, 1951). This identification was confirmed by searching BOLD with the two DNA barcode sequences (Table 1).

**Fig. 1.**
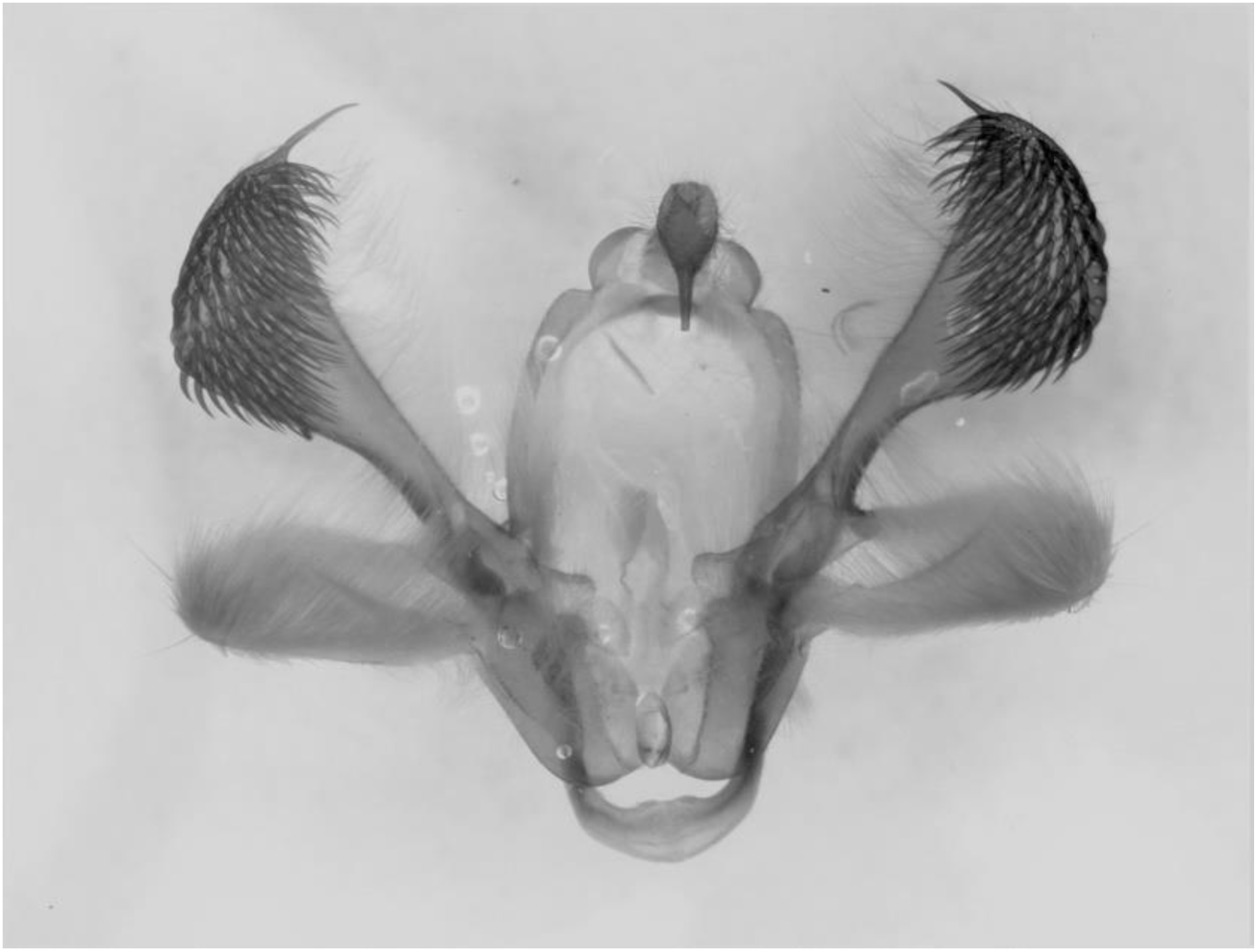
Valves of male genitalia of *Mythimna separata*, specimen am14016.

## Discussion

Initial attempts to identify our FAW samples using DNA barcodes were confounded by the existence of multiple species names for the two BINs (Barcode Index Numbers, essentially clusters of closely related sequences; Ratnasingham and Hebert, 2013) that comprise this species. The five species names recorded on BOLD at this time for *Spodoptera frugiperda* sequences are: *Spodoptera frugiperda, Spodoptera* “frugiperda sp. 1”, *Spodoptera* “frugiperda sp. 2,” *Spodoptera* “frugiperdaDHJ01” and *Spodoptera* “frugiperdaDHJ02”. In addition, there are other samples in the two BOLD BINs that have been identified only to genus level, or only to Order (Lepidoptera). When taken together with the multiple misidentifications on BOLD and GenBank (Table 3) and the lack of sequence quality control (Table 2), it was clear that in its current format, lacking adequate checks and balances, naive use of BOLD for identifying this species was misleading. Therefore, we have provided a dataset of vetted barcode sequences to be used for FAW identification.

COI barcode sequences from Bhutan samples are identical to some of those from both neighboring countries India and China, and from Myanmar and Vietnam. In addition to its origin in the Americas, this haplotype is also found in West, East and Southern Africa. Thus DNA barcode data does not help answer the question of the origin of the Bhutan incursion. However, based on the terrain of the country, with the Himalayan high ranges between Bhutan and China, it is most likely that they originated in India.

Fall Armyworm was first detected in Bhutan in September 2019, in the western district of Punakha, infesting maize. It was also detected in the southern district of Chhukha in December 2019, with subsequent infestation in maize in April 2020 in the same location. Given that the species is also present in neighboring countries, and that it is highly migratory (Nagoshi et al., 2012; Westbrook et al., 2016) and polyphagous, it is deemed established in Bhutan.

The FAW in the USA overwinters in the southern states of Texas and Florida and migrates to the northern states every year (Nagoshi et al., 2012). This is because the FAW is a pest of tropical origin unable to undergo diapause and cannot survive cold conditions as its development, reproduction and distribution is highly temperature dependent (Du Plessis et al., 2020; He et al., 2019; Sparks, 1980). In Bhutan, it may not survive the cold winters or be able to overwinter in the temperate regions such as the ones from which the samples for this study were collected. Nevertheless, FAW likely has established as a permanent, multigenerational pest in the southern region of the country which has sub-tropical climatic conditions from which it might migrate to the interior, cooler temperate regions of the country with the availability of host plants and onset of favourable climatic conditions.

Identifying whether an incursion is either rice or corn strain is important since the strains not only show different host plant preferences (the corn strain being more restricted to maize, while the rice strain is polyphagous) but also differ in many other biological characteristics, including susceptibility to different insecticides (Adamczyk et al., 1997; Ríos-Díez and Saldamando-Benjumea, 2011). However, Zhang et al. (2020) sequenced the genome of *S. frugiperda* and examined the population structure of the species in China, finding that it comprises a complex hybrid of rice and corn strains. This demonstrates that identifying strains using either the mitochondrial COI gene or the nuclear gene Triose phosphate isomerase (*Tpi*; Nagoshi, 2010) can be an oversimplification giving an inaccurate picture of the expected traits of an invading population.

The current recommended practice for management of FAW in Bhutan includes monitoring adult activity by deploying pheromone traps and with the capture of three adult moths per trap sprays with biopesticides such as neem oil is recommended. For maize, observed damage levels of 10-30% and 30-50% at the early and late whorl stages, respectively, will require pesticide applications. At the final growth stages, the tassel and silk stages, no pesticide spray is recommended, but mechanical control of the larval stages and conservation of natural enemy is advocated.

Northern Armyworm, *M. separata*, was described from Shanghai in 1865. It is distributed from Afghanistan eastwards through India, across Asia and south to Australia and New Zealand. It is listed by CABI as present in Nepal, but Bhutan is not listed, despite the country likely being part of its native range. In 2013, an outbreak of this species was recorded in eighteen districts of Bhutan attacking crops such as rice and maize. Initial infestation started on rice nurseries with subsequent, large scale infestation observed on maize across the country. The pest was however bought under control with sprays of organophosphates, pyrethroids and neem-based products. Timely sprays have been observed to reduce crop damage and average crop recovery rates between 30-70% in maize were observed after pesticide application.

## Conclusion

We have provided a reference DNA barcode dataset that can be used to definitively identify Fall Armyworm to species. Identifying corn and rice strains of FAW requires more than just a barcode sequence, however we note that some authors question the utility of the concept of corn and rice strains in this species. FAW is now present in Bhutan, and the presence of this new pest requires the establishment of a robust field monitoring, surveillance and scouting system along with programs aimed at raising growers’ knowledge in managing this pest. While interim management practices against FAW have been recommended, the biology and the occurrence of natural enemies of this pest species in Bhutan requires further investigation to develop an integrated pest management (IPM) strategies that is suitable for Bhutanese farmers. Northern Armyworm is now a major pest in all major rice and maize growing districts in Bhutan. For early detection and management, regular field surveillance and pheromone trapping is implemented.

## Supporting information

Dataset in Fasta format

## Acknowledgements

This research was funded by the Australian Museum Research Institute and the Department of Agriculture, Ministry of Agriculture and Forests, Bhutan.

## Supplementary data

Supplementary data to this article can be found online.

## References

Adamczyk, J.J.J., Holloway, J.W., Leonard, B.R., Graves, J.B., 1997. Susceptibility of fall armyworm collected from different plant hosts to selected insecticides and transgenic Bt cotton. Journal of Cotton Science 1, 21–28.

Bhavani, B., Chandra, S.V., Kishore, V.P., Bharatha, L.M., Jamuna, P., Swapna, B., 2019. Morphological and molecular identification of an invasive insect pest, fall army worm, Spodoptera frugiperda occurring on sugarcane in Andhra Pradesh, India. Journal of Entomology and Zoology Studies, 7(4): 12–18

CABI, 2020. Spodoptera frugiperda (fall armyworm). In: Invasive Species Compendium. Wallingford, UK: CAB International. www.cabi.org/isc (Accessed 21 April 2020).

Day, R., Abrahams, P., Bateman, M., Beale, T., Clottey, V., Cock, M., Colmenarez, Y., Corniani, N., Early., R, Godwin, J., Gomez, J., Moreno, P.G., Murphy, S.T., Oppong-Mensah, B., Phiri, N., Pratt, C., Silvestri, S., Witt, A., 2017. Fall armyworm: impacts and implications for Africa. Outlooks on Pest Management 28, 196–201.

Du Plessis, H., Schlemmer, M.-L., Van den Berg, J., 2020. The effect of temperature on the development of *Spodoptera frugiperda* (Lepidoptera: Noctuidae). Insects 11, 228.

Franclemont, J.G., 1951. The species of the *Leucania unipuncta* group, with a discussion of the generic names for the various segregates of *Leucania* in North America. Proceedings of the Entomological Society of Washington 53, 57–85.

Gilligan, T.M., Passoa, S.C., 2014. LepIntercept, an identification resource for intercepted Lepidoptera larvae. Identification Technology Program (ITP), USDA-APHIS-PPQ- S&T, Fort Collins, USA. (Accessed at www.lepintercept.org)

Goergen, G., Kumar P.L., Sankung, S.B., Togola, A., Tamò, M., 2016. First report of outbreaks of the fall armyworm *Spodoptera frugiperda* (J E Smith) (Lepidoptera, Noctuidae), a new alien invasive pest in west and central Africa. PLoS ONE 11, e0165632.

He, L.M., Ge, S.S., Chen, Y.C., Wu, Q.L., Jiang, Y. Y., Wu, K.M., 2019. The developmental threshold temperature, effective accumulated temperature and prediction model of developmental duration of fall armyworm, *Spodoptera frugiperda.* Plant Protection 45, 18–26.

Kalleshwaraswamy, C.M., Asokan, R., MahadevaSwamy, H.M., Marutid, M.S., Pavithra, H.B., Hegde, K., Navi, S., Prabhu, S.T., Goergen, G., 2018. First report of the fall armyworm, *Spodoptera frugiperda* (J. E. Smith) (Lepidoptera: Noctuidae), an alien invasive pest on maize in India. Pest Management in Horticultural Ecosystems 4, 23–29.

Kearse, M., Moir, R., Wilson, A., Stones-Havas, S., Cheung, M., Sturrock, S., Buxton, S., Cooper, A., Markowitz, S., Duran, C., Thierer, T., Ashton, B., Mentjies, P., Drummond, A., 2012. Geneious Basic: an integrated and extendable desktop software platform for the organization and analysis of sequence data. Bioinformatics 28, 1647–1649.

Lee, T.R.C., Anderson, S.J., Tran-Nguyen, L.T.T., Sallam, N., Le Ru, B.P., Conlong, D., Powell, K., Ward, A., Mitchell, A., 2019. Towards a global DNA barcode reference library for quarantine identifications of lepidopteran stemborers, with an emphasis on sugarcane pests. Scientific Reports 9, 7039.

Mitchell, A., 2015. Collecting in collections: a PCR strategy and primer set for DNA barcoding of decades-old dried museum specimens. Molecular Ecology Resources 15, 1102–1111.

Mitchell, A., Gopurenko, D., 2016. DNA barcoding the Heliothinae (Lepidoptera: Noctuidae) of Australia and utility of DNA barcodes for pest identiifcation in *Helicoverpa* and relatives. PLoS ONE 11, e0160895.

Montezano, D.G., Specht, A., Sosa-Gomez, D.R., Roque-Specht, V.F., Sousa-Silva, J.C., Paula-Moraes, S.V., Peterson, J.A., Hunt, T.E., 2018. Host plants of *Spodoptera frugiperda* (Lepidoptera: Noctuidae) in the Americas. African Entomology 26, 286–301.

Nagoshi, R.N., 2010. The fall armyworm triose phosphate isomerase (*Tpi*) gene as a marker of strain identity and interstrain mating. Annals of the Entomological Society of America 103(2), 283–292.

Nagoshi, R.N., Meagher, R.L., Hay-Roe, M., 2012. Inferring the annual migration patterns of fall armyworm (Lepidoptera: Noctuidae) in the United States from mitochondrial haplotypes. Ecology and Evolution 2, 1458–67.

Pogue, M.G., 2002. A world revision of the genus *Spodoptera* Guenée (Lepidoptera: Noctuidae). Memoirs of the American Entomological Society 43, 1–201.

Ratnasingham, S., Hebert, P.D.N., 2013. A DNA-Based Registry for All Animal Species: The Barcode Index Number (BIN) System. PLoS ONE 8, e66213.

Ríos-Díez, J.D., Saldamando-Benjumea, C.I., 2011. Susceptibility of *Spodoptera frugiperda* (Lepidoptera: Noctuidae) strains from central Colombia to two insecticides, methomyl and lambda-cyhalothrin: A study of the genetic basis of resistance, Journal of Economic Entomology 104, 1698–1705.

Sparks, A.N., 1980. A review of the biology of the fall armyworm. Florida Entomology 62, 82–87.

Villesen, P., 2007. FaBox: an online toolbox for FASTA sequences. Molecular Ecology Notes 7, 965–968.

Westbrook, J.K., Nagoshi, R.N., Meagher, R.L., Fleischer, S.J., Jairam, S., 2016. Modeling seasonal migration of fall armyworm moths. International Journal of Biometeorology 60, 255–267.

Zhang, L., Liu, B., Zheng, W., Liu, C., Zhang, D., Zhao, S., Xu, P., Wilson, K., Withers, A., Jones, C.M., Smith, J.A., Chipabika, G., Kachigamba, G.L., Nam, K., d’Alençon, E., Liu, B., Liang, X., Jin, M., Wu, C., Chakrabarty, S., Yang, X., Jiang, Y., Liu, J., Liu, X., Quan, W., Wang, G., Fan, W., Qian, W., Wu, K., Xiao, Y., 2020 (Preprint). High-depth resequencing reveals hybrid population and insecticide resistance characteristics of fall armyworm (*Spodoptera frugiperda*) invading China. Biorxiv doi: https://doi.org/10.1101/813154

