## Supplementary Figure S1 for "An updated global COI barcode reference data set for Fall Armyworm (*Spodoptera frugiperda*) and first record of this species in Bhutan"

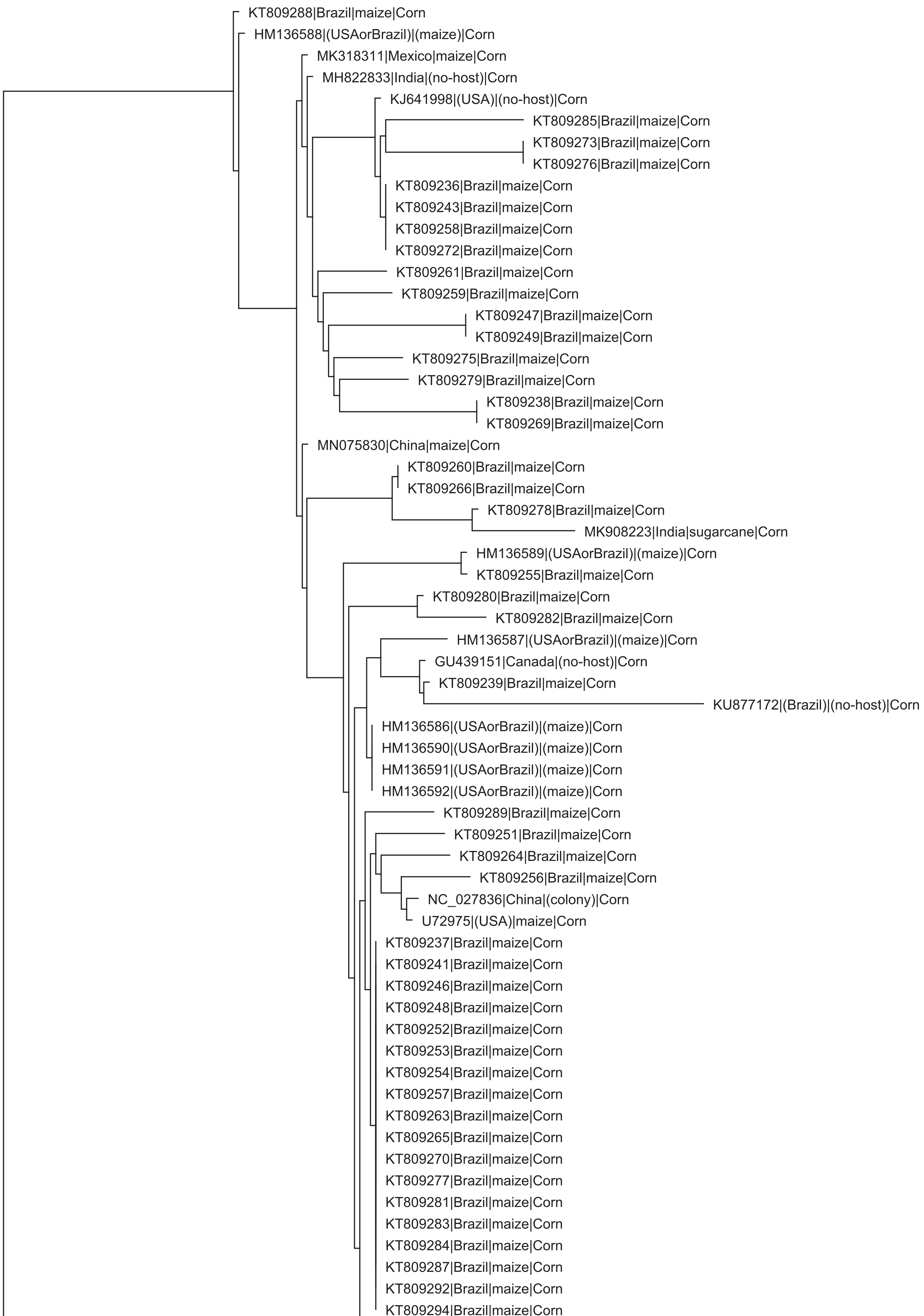

KT809288|Brazil|maize|Corn
HM136588|(USAorBrazil)|(maize)|Corn
MK318311|Mexico|maize|Corn
MH822833|India|(no-host)|Corn
KJ641998|(USA)|(no-host)|Corn
KT809285|Brazil|maize|Corn
KT809273|Brazil|maize|Corn
KT809276|Brazil|maize|Corn
KT809236|Brazil|maize|Corn
KT809243|Brazil|maize|Corn
KT809258|Brazil|maize|Corn
KT809272|Brazil|maize|Corn
KT809261|Brazil|maize|Corn
KT809259|Brazil|maize|Corn
KT809247|Brazil|maize|Corn
KT809249|Brazil|maize|Corn
KT809275|Brazil|maize|Corn
KT809279|Brazil|maize|Corn
KT809238|Brazil|maize|Corn
KT809269|Brazil|maize|Corn
MN075830|China|maize|Corn
KT809260|Brazil|maize|Corn
KT809266|Brazil|maize|Corn
KT809278|Brazil|maize|Corn
MK908223|India|sugarcane|Corn
HM136589|(USAorBrazil)|(maize)|Corn
KT809255|Brazil|maize|Corn
KT809280|Brazil|maize|Corn
KT809282|Brazil|maize|Corn
HM136587|(USAorBrazil)|(maize)|Corn
GU439151|Canada|(no-host)|Corn
KT809239|Brazil|maize|Corn
KU877172|(Brazil)|(no-host)|Corn
HM136586|(USAorBrazil)|(maize)|Corn
HM136590|(USAorBrazil)|(maize)|Corn
HM136591|(USAorBrazil)|(maize)|Corn
HM136592|(USAorBrazil)|(maize)|Corn
KT809289|Brazil|maize|Corn
KT809251|Brazil|maize|Corn
KT809264|Brazil|maize|Corn
KT809256|Brazil|maize|Corn
NC_027836|China|(colony)|Corn
U72975|(USA)|maize|Corn
KT809237|Brazil|maize|Corn
KT809241|Brazil|maize|Corn
KT809246|Brazil|maize|Corn
KT809248|Brazil|maize|Corn
KT809252|Brazil|maize|Corn
KT809253|Brazil|maize|Corn
KT809254|Brazil|maize|Corn
KT809257|Brazil|maize|Corn
KT809263|Brazil|maize|Corn
KT809265|Brazil|maize|Corn
KT809270|Brazil|maize|Corn
KT809277|Brazil|maize|Corn
KT809281|Brazil|maize|Corn
KT809283|Brazil|maize|Corn
KT809284|Brazil|maize|Corn
KT809287|Brazil|maize|Corn
KT809292|Brazil|maize|Corn
KT809294|Brazil|maize|Corn

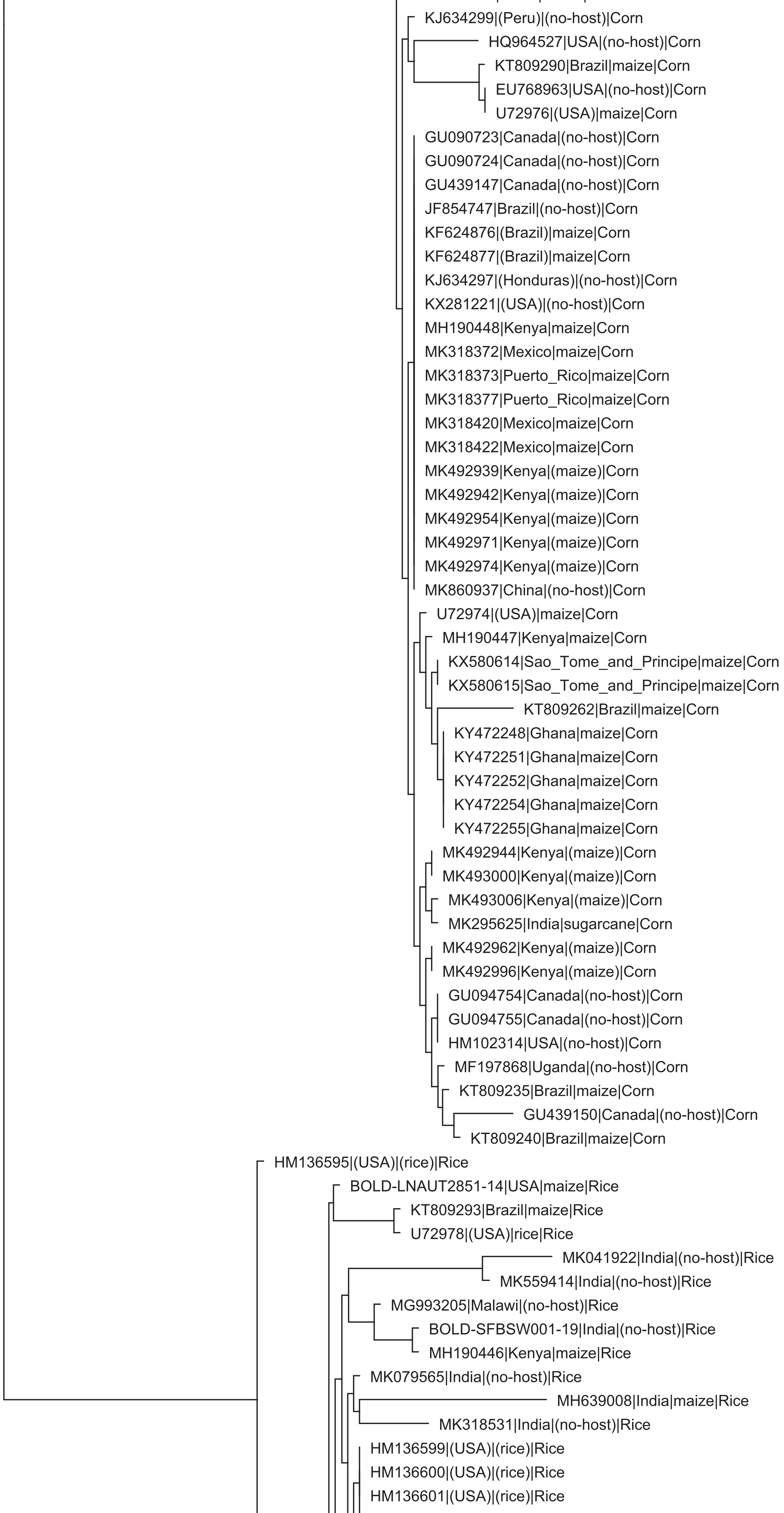

KJ634299|(Peru)|(no-host)|Corn
HQ964527|USA|(no-host)|Corn
KT809290|Brazil|maize|Corn
EU768963|USA|(no-host)|Corn
U72976|(USA)|maize|Corn
GU090723|Canada|(no-host)|Corn
GU090724|Canada|(no-host)|Corn
GU439147|Canada|(no-host)|Corn
JF854747|Brazil|(no-host)|Corn
KF624876|(Brazil)|maize|Corn
KF624877|(Brazil)|maize|Corn
KJ634297|(Honduras)|(no-host)|Corn
KX281221|(USA)|(no-host)|Corn
MH190448|Kenya|maize|Corn
MK318372|Mexico|maize|Corn
MK318373|Puerto_Rico|maize|Corn
MK318377|Puerto_Rico|maize|Corn
MK318420|Mexico|maize|Corn
MK318422|Mexico|maize|Corn
MK492939|Kenya|(maize)|Corn
MK492942|Kenya|(maize)|Corn
MK492954|Kenya|(maize)|Corn
MK492971|Kenya|(maize)|Corn
MK492974|Kenya|(maize)|Corn
MK860937|China|(no-host)|Corn
U72974|(USA)|maize|Corn
MH190447|Kenya|maize|Corn
KX580614|Sao_Tome_and_Principe|maize|Corn
KX580615|Sao_Tome_and_Principe|maize|Corn
KT809262|Brazil|maize|Corn
KY472248|Ghana|maize|Corn
KY472251|Ghana|maize|Corn
KY472252|Ghana|maize|Corn
KY472254|Ghana|maize|Corn
KY472255|Ghana|maize|Corn
MK492944|Kenya|(maize)|Corn
MK493000|Kenya|(maize)|Corn
MK493006|Kenya|(maize)|Corn
MK295625|India|sugarcane|Corn
MK492962|Kenya|(maize)|Corn
MK492996|Kenya|(maize)|Corn
GU094754|Canada|(no-host)|Corn
GU094755|Canada|(no-host)|Corn
HM102314|USA|(no-host)|Corn
MF197868|Uganda|(no-host)|Corn
KT809235|Brazil|maize|Corn
GU439150|Canada|(no-host)|Corn
KT809240|Brazil|maize|Corn
HM136595|(USA)|(rice)|Rice
BOLD-LNAUT2851-14|USA|maize|Rice
KT809293|Brazil|maize|Rice
U72978|(USA)|rice|Rice
MK041922|India|(no-host)|Rice
MK559414|India|(no-host)|Rice
MG993205|Malawi|(no-host)|Rice
BOLD-SFBSW001-19|India|(no-host)|Rice
MH190446|Kenya|maize|Rice
MK079565|India|(no-host)|Rice
MH639008|India|maize|Rice
MK318531|India|(no-host)|Rice
HM136599|(USA)|(rice)|Rice
HM136600|(USA)|(rice)|Rice
HM136601|(USA)|(rice)|Rice

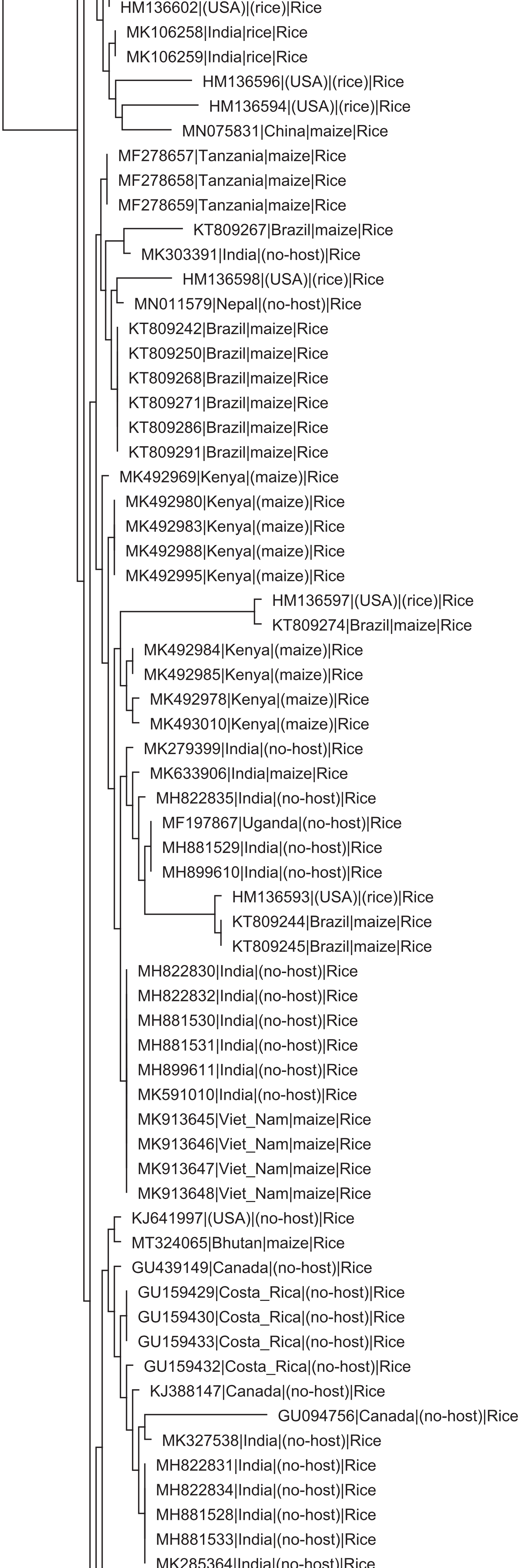
HM136602|(USA)|(rice)|Rice
MK106258|India|rice|Rice
MK106259|India|rice|Rice
HM136596|(USA)|(rice)|Rice
HM136594|(USA)|(rice)|Rice
MN075831|China|maize|Rice
MF278657|Tanzania|maize|Rice
MF278658|Tanzania|maize|Rice
MF278659|Tanzania|maize|Rice
KT809267|Brazil|maize|Rice
MK303391|India|(no-host)|Rice
HM136598|(USA)|(rice)|Rice
MN011579|Nepal|(no-host)|Rice
KT809242|Brazil|maize|Rice
KT809250|Brazil|maize|Rice
KT809268|Brazil|maize|Rice
KT809271|Brazil|maize|Rice
KT809286|Brazil|maize|Rice
KT809291|Brazil|maize|Rice
MK492969|Kenya|(maize)|Rice
MK492980|Kenya|(maize)|Rice
MK492983|Kenya|(maize)|Rice
MK492988|Kenya|(maize)|Rice
MK492995|Kenya|(maize)|Rice
HM136597|(USA)|(rice)|Rice
KT809274|Brazil|maize|Rice
MK492984|Kenya|(maize)|Rice
MK492985|Kenya|(maize)|Rice
MK492978|Kenya|(maize)|Rice
MK493010|Kenya|(maize)|Rice
MK279399|India|(no-host)|Rice
MK633906|India|maize|Rice
MH822835|India|(no-host)|Rice
MF197867|Uganda|(no-host)|Rice
MH881529|India|(no-host)|Rice
MH899610|India|(no-host)|Rice
HM136593|(USA)|(rice)|Rice
KT809244|Brazil|maize|Rice
KT809245|Brazil|maize|Rice
MH822830|India|(no-host)|Rice
MH822832|India|(no-host)|Rice
MH881530|India|(no-host)|Rice
MH881531|India|(no-host)|Rice
MH899611|India|(no-host)|Rice
MK591010|India|(no-host)|Rice
MK913645|Viet_Nam|maize|Rice
MK913646|Viet_Nam|maize|Rice
MK913647|Viet_Nam|maize|Rice
MK913648|Viet_Nam|maize|Rice
KJ641997|(USA)|(no-host)|Rice
MT324065|Bhutan|maize|Rice
GU439149|Canada|(no-host)|Rice
GU159429|Costa_Rica|(no-host)|Rice
GU159430|Costa_Rica|(no-host)|Rice
GU159433|Costa_Rica|(no-host)|Rice
GU159432|Costa_Rica|(no-host)|Rice
KJ388147|Canada|(no-host)|Rice
GU094756|Canada|(no-host)|Rice
MK327538|India|(no-host)|Rice
MH822831|India|(no-host)|Rice
MH822834|India|(no-host)|Rice
MH881528|India|(no-host)|Rice
MH881533|India|(no-host)|Rice
MK285364|India|(no-host)|Rice

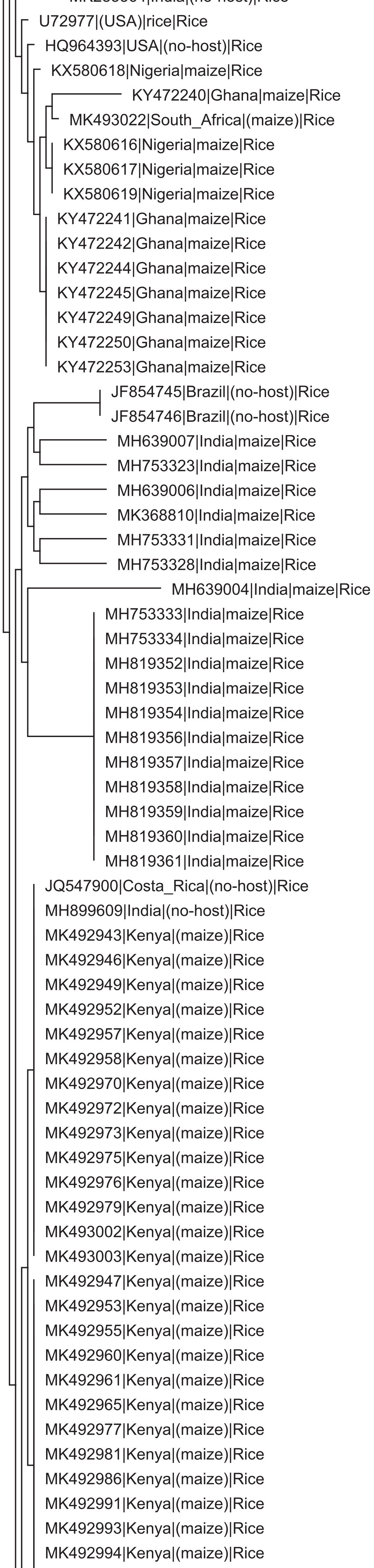
U72977|(USA)|rice|Rice
HQ964393|USA|(no-host)|Rice
KX580618|Nigeria|maize|Rice
KY472240|Ghana|maize|Rice
MK493022|South_Africa|(maize)|Rice
KX580616|Nigeria|maize|Rice
KX580617|Nigeria|maize|Rice
KX580619|Nigeria|maize|Rice
KY472241|Ghana|maize|Rice
KY472242|Ghana|maize|Rice
KY472244|Ghana|maize|Rice
KY472245|Ghana|maize|Rice
KY472249|Ghana|maize|Rice
KY472250|Ghana|maize|Rice
KY472253|Ghana|maize|Rice
JF854745|Brazil|(no-host)|Rice
JF854746|Brazil|(no-host)|Rice
MH639007|India|maize|Rice
MH753323|India|maize|Rice
MH639006|India|maize|Rice
MK368810|India|maize|Rice
MH753331|India|maize|Rice
MH753328|India|maize|Rice
MH639004|India|maize|Rice
MH753333|India|maize|Rice
MH753334|India|maize|Rice
MH819352|India|maize|Rice
MH819353|India|maize|Rice
MH819354|India|maize|Rice
MH819356|India|maize|Rice
MH819357|India|maize|Rice
MH819358|India|maize|Rice
MH819359|India|maize|Rice
MH819360|India|maize|Rice
MH819361|India|maize|Rice
JQ547900|Costa_Rica|(no-host)|Rice
MH899609|India|(no-host)|Rice
MK492943|Kenya|(maize)|Rice
MK492946|Kenya|(maize)|Rice
MK492949|Kenya|(maize)|Rice
MK492952|Kenya|(maize)|Rice
MK492957|Kenya|(maize)|Rice
MK492958|Kenya|(maize)|Rice
MK492970|Kenya|(maize)|Rice
MK492972|Kenya|(maize)|Rice
MK492973|Kenya|(maize)|Rice
MK492975|Kenya|(maize)|Rice
MK492976|Kenya|(maize)|Rice
MK492979|Kenya|(maize)|Rice
MK493002|Kenya|(maize)|Rice
MK493003|Kenya|(maize)|Rice
MK492947|Kenya|(maize)|Rice
MK492953|Kenya|(maize)|Rice
MK492955|Kenya|(maize)|Rice
MK492960|Kenya|(maize)|Rice
MK492961|Kenya|(maize)|Rice
MK492965|Kenya|(maize)|Rice
MK492977|Kenya|(maize)|Rice
MK492981|Kenya|(maize)|Rice
MK492986|Kenya|(maize)|Rice
MK492991|Kenya|(maize)|Rice
MK492993|Kenya|(maize)|Rice
MK492994|Kenya|(maize)|Rice

MK492998|Kenya|(maize)|Rice
MK492999|Kenya|(maize)|Rice
MK493004|Kenya|(maize)|Rice
MK493008|Kenya|(maize)|Rice
BOLD-AGIMP054-18|India|(no-host)|Rice
BOLD-FAW001-19|India|maize|Rice
GU095403|Canada|(no-host)|Rice
GU159426|Costa_Rica|(no-host)|Rice
GU159427|Costa_Rica|(no-host)|Rice
GU159428|Costa_Rica|(no-host)|Rice
GU159431|Costa_Rica|(no-host)|Rice
GU159434|Costa_Rica|(no-host)|Rice
GU159435|Costa_Rica|(no-host)|Rice
GU163698|Costa_Rica|(no-host)|Rice
GU439148|Canada|(no-host)|Rice
HQ964351|USA|(no-host)|Rice
HQ964352|USA|(no-host)|Rice
HQ964353|USA|(no-host)|Rice
HQ964394|USA|(no-host)|Rice
HQ964439|USA|(no-host)|Rice
HQ964440|USA|(no-host)|Rice
HQ964441|USA|(no-host)|Rice
HQ964442|USA|(no-host)|Rice
HQ964443|USA|(no-host)|Rice
HQ964485|USA|(no-host)|Rice
HQ964486|USA|(no-host)|Rice
HQ964487|USA|(no-host)|Rice
JF854740|Brazil|(no-host)|Rice
JF854741|Brazil|(no-host)|Rice
JF854743|Brazil|(no-host)|Rice
JF854744|Brazil|(no-host)|Rice
JQ554012|Costa_Rica|(no-host)|Rice
JQ559528|Costa_Rica|(no-host)|Rice
JQ571459|Costa_Rica|(no-host)|Rice
JQ572603|Costa_Rica|(no-host)|Rice
JQ577923|Costa_Rica|(no-host)|Rice
KJ634298|(Suriname)|(no-host)|Rice
MH190444|Kenya|maize|Rice
MH190445|Kenya|maize|Rice
MH639005|India|maize|Rice
MH704433|India|(no-host)|Rice
MH753324|India|sorghum|Rice
MH753325|India|maize|Rice
MH753326|India|maize|Rice
MH753327|India|maize|Rice
MH753329|India|maize|Rice
MH753330|India|maize|Rice
MH753332|India|maize|Rice
MK105749|India|rice|Rice
MK105750|India|rice|Rice
MK318297|Dominican_Republic|Capsicum|Rice
MK492929|Kenya|(maize)|Rice
MK492930|Kenya|(maize)|Rice
MK492931|Kenya|(maize)|Rice
MK492932|Kenya|(maize)|Rice
MK492933|Kenya|(maize)|Rice
MK492934|Kenya|(maize)|Rice
MK492935|Kenya|(maize)|Rice
MK492936|Kenya|(maize)|Rice
MK492937|Kenya|(maize)|Rice
MK492938|Kenya|(maize)|Rice
MK492940|Kenya|(maize)|Rice
MK492941|Kenya|(maize)|Rice
MK492945|Kenya|(maize)|Rice

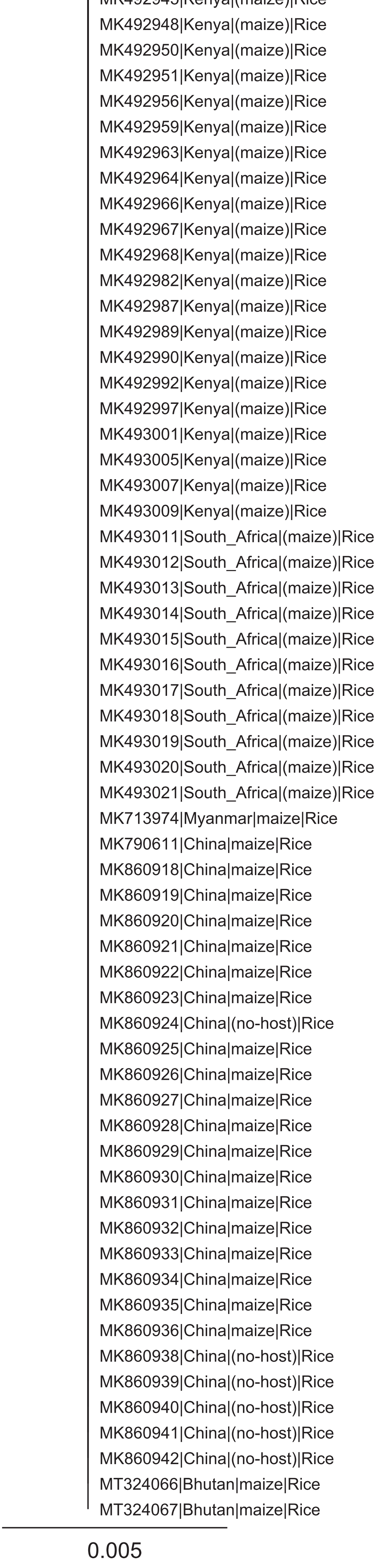
MK492948|Kenya|(maize)|Rice
MK492950|Kenya|(maize)|Rice
MK492951|Kenya|(maize)|Rice
MK492956|Kenya|(maize)|Rice
MK492959|Kenya|(maize)|Rice
MK492963|Kenya|(maize)|Rice
MK492964|Kenya|(maize)|Rice
MK492966|Kenya|(maize)|Rice
MK492967|Kenya|(maize)|Rice
MK492968|Kenya|(maize)|Rice
MK492982|Kenya|(maize)|Rice
MK492987|Kenya|(maize)|Rice
MK492989|Kenya|(maize)|Rice
MK492990|Kenya|(maize)|Rice
MK492992|Kenya|(maize)|Rice
MK492997|Kenya|(maize)|Rice
MK493001|Kenya|(maize)|Rice
MK493005|Kenya|(maize)|Rice
MK493007|Kenya|(maize)|Rice
MK493009|Kenya|(maize)|Rice
MK493011|South_Africa|(maize)|Rice
MK493012|South_Africa|(maize)|Rice
MK493013|South_Africa|(maize)|Rice
MK493014|South_Africa|(maize)|Rice
MK493015|South_Africa|(maize)|Rice
MK493016|South_Africa|(maize)|Rice
MK493017|South_Africa|(maize)|Rice
MK493018|South_Africa|(maize)|Rice
MK493019|South_Africa|(maize)|Rice
MK493020|South_Africa|(maize)|Rice
MK493021|South_Africa|(maize)|Rice
MK713974|Myanmar|maize|Rice
MK790611|China|maize|Rice
MK860918|China|maize|Rice
MK860919|China|maize|Rice
MK860920|China|maize|Rice
MK860921|China|maize|Rice
MK860922|China|maize|Rice
MK860923|China|maize|Rice
MK860924|China|(no-host)|Rice
MK860925|China|maize|Rice
MK860926|China|maize|Rice
MK860927|China|maize|Rice
MK860928|China|maize|Rice
MK860929|China|maize|Rice
MK860930|China|maize|Rice
MK860931|China|maize|Rice
MK860932|China|maize|Rice
MK860933|China|maize|Rice
MK860934|China|maize|Rice
MK860935|China|maize|Rice
MK860936|China|maize|Rice
MK860938|China|(no-host)|Rice
MK860939|China|(no-host)|Rice
MK860940|China|(no-host)|Rice
MK860941|China|(no-host)|Rice
MK860942|China|(no-host)|Rice
MT324066|Bhutan|maize|Rice
MT324067|Bhutan|maize|Rice
0.005
